# DecoPath: A web application for decoding pathway enrichment analysis

**DOI:** 10.1101/2021.05.22.445243

**Authors:** Sarah Mubeen, Vinay Srinivas Bharadhwaj, Yojana Gadiya, Martin Hofmann-Apitius, Alpha Tom Kodamullil, Daniel Domingo-Fernández

## Abstract

The past two decades have brought a steady growth of pathway databases and pathway enrichment methods. However, the advent of pathway data has not been accompanied by an improvement with regards to interoperability across databases, thus, hampering the use of pathway knowledge from multiple databases for pathway enrichment analyses. While integrative databases have attempted to address this issue by collating pathway knowledge from multiple resources, these approaches do not account for redundant information across them. On the other hand, the majority of studies that employ pathway enrichment analyses still rely upon a single database, though the use of another resource could yield differing results, which is similarly the case when different pathway enrichment methods are employed. These shortcomings call for approaches that investigate the differences and agreements across databases and enrichment methods as their selection in the experimental design of a pathway analysis can be a crucial first step in ensuring the results of such an analysis are meaningful. Here we present DecoPath, a web application to assist in the interpretation of the results of pathway enrichment analysis. DecoPath provides an ecosystem to run pathway enrichment analysis or directly upload results and facilitate the interpretation of these results with custom visualizations that highlight the consensus and/or discrepancies at the pathway- and gene-levels. DecoPath is available at https://decopath.scai.fraunhofer.de and its source code and documentation can be found on GitHub at https://github.com/DecoPath/DecoPath.

## 1. Introduction

In recent years, high throughput (HT) technologies have given rise to a perpetual influx of *-omics* data, requiring pragmatic approaches to sift out meaning. One of the most common applications of HT technologies is gene expression profiling to simultaneously determine the expression patterns of thousands of genes at the transcription level under certain conditions (Dillies *et al*., 2013). While a host of statistical techniques are available to identify genes that differ in expression depending on a particular condition, gene set or pathway enrichment analysis methods represent a major class of tools researchers employ to group lists of genes into defined pathways and understand the functional roles of genes for any given set of conditions (Reimand *et al*., 2019). To date, almost a hundred different pathway enrichment methods have been proposed, including the popular over-representation analysis (ORA) and gene set enrichment analysis (GSEA) (Subramanian *et al*., 2005). Though these methods may vary based on the overarching categories they fall into (e.g., topology vs. non-topology-based) or the statistical techniques used, they have widely shown their ability to deconvolute biological pathways dysregulated in a given state (Nguyen *et al*., 2019).

Numerous pathway databases have been developed which aim at representing biological pathways from various vantage points (e.g., differing scopes, contexts, boundaries or pathway types). The existence of several hundreds of these databases reflects the inherent complexity and variability of biological processes that occur in living organisms. Further compounding this complexity is the fact that biological pathways housed in these databases are human constructs, delimited based on abstract boundaries defined by a researcher or the consensus of the community. This implies that a well-studied pathway could contain different biological entities depending on the boundaries defined by the databases that store it. These differences across databases can manifest in variability in the results of pathway enrichment analysis depending on both the method (Geistlinger *et al*., 2020; Nguyen *et al*., 2019; Zyla *et al*., 2019; Mathur *et al*., 2018) as well as the pathway database employed (Mubeen *et al*., 2019).

Recent approaches to pathway enrichment analysis have focused on the integration of multiple datasets across different platforms to ensure a broader coverage of significantly enriched pathways (Griss *et al*., 2020; Paczkowska *et al*., 2020; Zhou *et al*., 2019). Other techniques attempt to account for potential differences that may arise in the results of pathway enrichment analysis by combining gene sets from several pathway databases. For instance, Canzler and Hackermüller (2020) presented an approach that leverages GSEA to calculate a combined enrichment score for multiple -*omics* layers using several databases. However, performing pathway enrichment analysis using multiple databases to increase the number of pathways covered can only partially address the challenges associated with variability in results. This is because such an approach falls short of leveraging the substantial overlap of pathway knowledge across databases which could provide more comprehensive results (Stobbe *et al*., 2011; Belenky *et al*., 2015; Domingo-Fernández *et al*., 2018). Furthermore, combining several databases can result in redundant pathways, an issue tackled by the SetRank algorithm which discounts significant gene sets if their significance can be explained by their overlap with another gene set (Simillion *et al*., 2017). Finally, a possible, natural solution to better connect and structure redundant information across databases lies in leveraging pathway ontologies (Petri *et al*., 2014) or pathway mappings with database cross-references (Domingo-Fernández *et al*. 2018). By connecting related pathways across databases, we can, in turn, investigate the consensus, or lack thereof, of the results of pathway enrichment analysis between databases or methods as demonstrated by several recent benchmarks (Geistlinger *et al*., 2020; Nguyen *et al*., 2019; Zyla *et al*., 2019; Mathur *et al*., 2018; Mubeen *et al*., 2019).

Here, we present DecoPath, a web application that provides a user-friendly and interactive application to compare and interpret the results of pathway enrichment analysis yielded by different pathway databases. To facilitate the comparison of results across databases, and bring to light possible contradictory results, we present several interactive visualization tools designed to better interpret the results of pathway enrichment at both the pathway and gene-level. While these visualizations can generally be used for any pathway enrichment method, DecoPath also integrates standard pathway enrichment methods in its pipeline, thus, enabling users to conduct an entire enrichment analysis on the web application (from data submission to interpretation). Finally, although DecoPath provides four default databases, it also allows users to upload gene sets and mappings such that analyses can be run on their independently curated gene sets.

## 2. Methods

### 2.1. Implementation

The server-side was implemented in the Python programming language using the Django framework (https://www.djangoproject.com/). This framework operates using a Model-View-Controller (MVC) architecture and was integrated with Celery (http://www.celeryproject.org) and RabbitMQ (https://www.rabbitmq.com) for asynchronous task execution. The front-end of DecoPath comprises several interactive visualizations implemented using a collection of powerful Javascript libraries, including jQuery (https://jquery.com), D3.js (https://d3js.org/), and DataTables (https://datatables.net/). Furthermore, DecoPath relies on Bootstrap 4 (https://getbootstrap.com/) for the main design of the website. The web application is containerized using Docker for reproducibility purposes and easy deployment.

### 2.2. Pathway resources

DecoPath enables users to compare the results of enrichment analysis yielded using various pathway databases. As mentioned in the Introduction, pathways in different databases can substantially overlap, such that a pathway in one database can have counterparts in several others. Leveraging equivalent pathway mappings across several widely-used databases, DecoPath aims at highlighting the consensus, or lack thereof, of enrichment analysis results for each equivalent pathway. Expanding upon our previous work (Domingo-Fernández *et al*., 2018), we added novel equivalent pathway mappings as well as mappings for an additional database (i.e., PathBank (Wishart *et al*., 2020)) **(Supplementary Text)**. Thus, the released version of DecoPath provides users with the following pathway databases: KEGG (Kanehisa *et al*., 2021), Reactome (Fabregat *et al*., 2018), WikiPathways (Martens *et al*., 2021), and PathBank (Retrieved 03.08.2020). Additionally, a DecoPath-specific gene set database containing merged gene sets of equivalent pathways across the aforementioned databases is also provided, as described in the following section.

### 2.3. Generating a pathway hierarchy

The consolidation of each of the pathway databases into a pathway meta-database was conducted in order to generate a pathway hierarchy. In doing so, equivalent representations of pathways across KEGG, PathBank, Reactome and WikiPathways were combined. The pathway hierarchy contains a total of 644 pathways from these four databases and can be found at https://github.com/ComPath/compath-resources/blob/master/mappings/decopath_ontology.xlsx (dated 01-13-2021). The hierarchy comprises seven major categories: metabolism, immune, signaling, communication and transport, cell-death, disease, DNA repair and replication, and others. All pathways in the hierarchy retained their original identifiers except equivalent pathways which were merged and given unique names and identifiers. The pathway hierarchy is a directed acyclic graph with a maximum depth of 4, in which relation types between pathways can be either *is-part-of* or *equivalent-to* relations. The curation process to generate the hierarchy is described in the **Supplementary Text**.

### 2.4. Pathway Enrichment Methods

DecoPath comprises two of the most widely used pathway enrichment methods (García-Campos *et al*., 2015; Katri *et al*., 2010; Xie *et al*., 2021): Over Representation Analysis (ORA) and Gene Set Enrichment Analysis (GSEA) (Subramanian *et al*., 2005). ORA aims at identifying pathways (i.e., gene sets) that are over-represented within a list of genes of interest. A pathway is considered enriched (over-represented) if the *p*-value arising from a one-sided Fisher’s exact test (Fisher, 1992) is lower than a specified threshold, typically 0.05. As this test is conducted for each pathway in the database, DecoPath’s implementation of ORA corrects the *p*-value by applying multiple hypothesis testing correction with the Benjamini–Yekutieli method under dependency (Benjamini and Yekutieli, 2001). The second method, GSEA, determines whether a pathway or a gene set significantly differs between two groups. A pathway is considered significantly regulated in that condition if genes of that pathway appear in the top or bottom ranking of a list of differentially expressed genes more than expected by chance. DecoPath uses the GSEA implementation from *gseapy* (https://gseapy.readthedocs.io/en/latest). Additionally, DecoPath enables conducting differential gene expression (DGE) analysis between groups through DESeq2 (version 1.22.2). Apart from these methods, DecoPath also provides the option to include additional pathway enrichment methods into the web application.

### 2.5. Installation

Although we provide a freely available instance of DecoPath at https://decopath.scai.fraunhofer.de/, in the case of large datasets or cases where the compute capacity of the server may be insufficient depending on the type of analysis, users can install and use DecoPath in their own system. We offer two options to install DecoPath depending on the needs of the user. The first and easiest method for those unfamiliar with Django-based web applications is to install Docker and deploy the Docker container which will install required components and run the web application. The second option is to directly deploy it following the instructions in the GitHub repository (https://github.com/decopath/decopath).

### 2.6. Run time considerations

Computation time is dependent on the type of analysis, size of the datasets as well as the device specifications. ORA can be run on a gene list on a timescale of seconds and requires the relatively lowest usage of memory. A DGE analysis task has a timescale of several minutes, while GSEA on a typical expression dataset with two experimental groups and four databases can also be done within minutes with a dual-core Intel Core i5 CPU and 16 GB RAM.

### 2.7. Case scenario

Using each of the available enrichment methods, we demonstrate a typical workflow in DecoPath with the The Cancer Genome Atlas Liver Hepatocellular Carcinoma (TCGA-LIHC) dataset (Weinstein *et al*., 2013). Gene expression data from this dataset was retrieved from the Genomic Data Commons (GDC; https://gdc.cancer.gov) portal through the R/Bioconductor package, TCGAbiolinks (version 2.16.3; [43]) on 04-08-2020. To run GSEA, we employed RNA-Seq expression data normalized using Fragments Per Kilobase of transcript per Million mapped reads upper quartile (FPKM-UQ). DGE analysis using read counts from the TCGA-LIHC dataset (retrieved from the GDC; https://gdc.cancer.gov) was performed between normal and tumor samples to derive a gene list to conduct ORA. This final list of genes was restricted to genes that exhibited an adjusted *p*-value < 0.05. Specifications of the parameter settings for ORA and GSEA are listed in **Supplementary Table 1**.

## 3. Results

Here, we describe the DecoPath web application. A typical workflow of the web application involves the submission of an experiment, generation of results, and the subsequent exploration and visualization of these results **(Figure 1)**. In the following, we provide a detailed description for each of the steps in the workflow.

**Figure 1.**
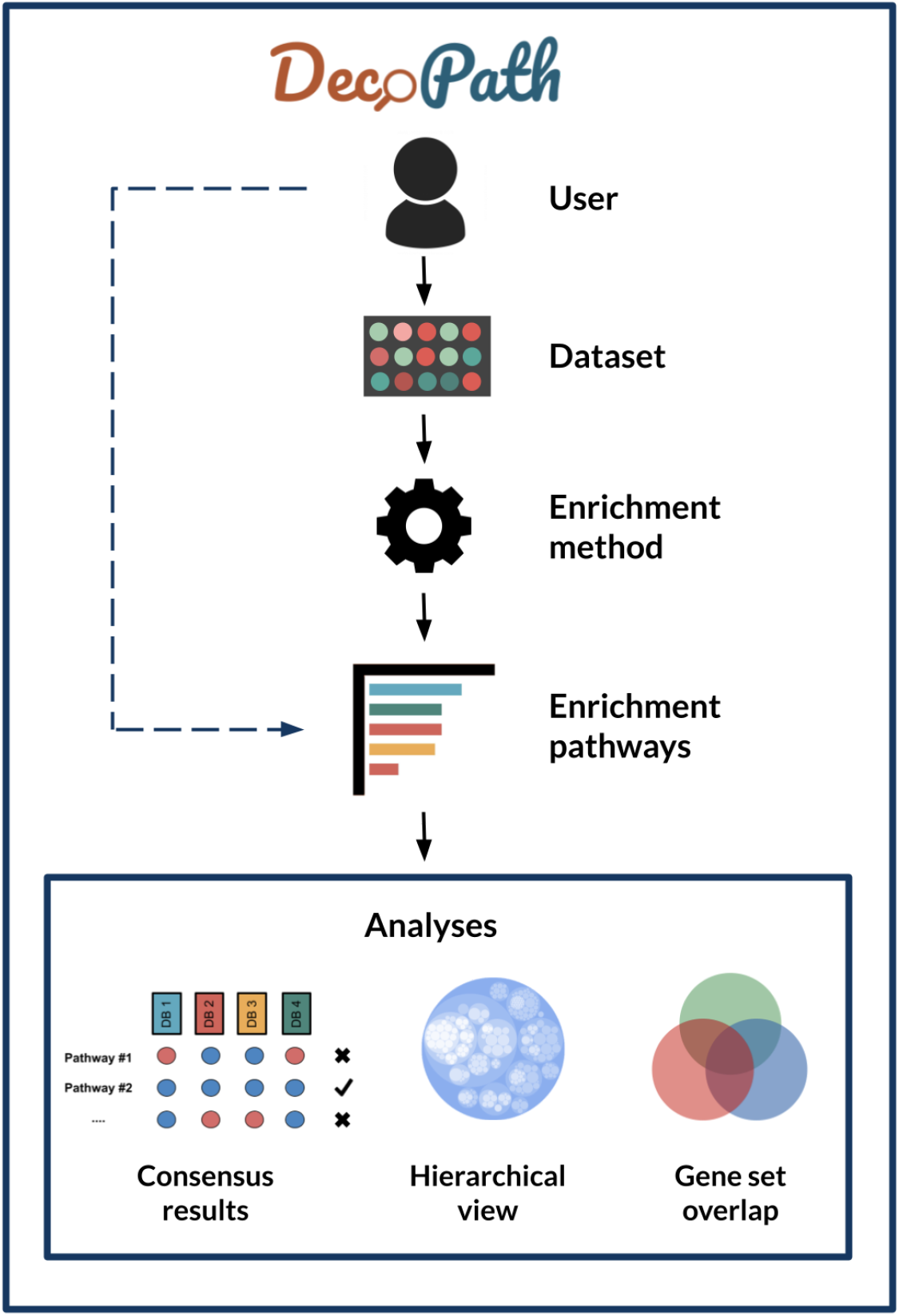
DecoPath workflow. Users can upload datasets to run pathway enrichment analysis or directly upload enrichment results from their own experiments. Once results have been loaded, DecoPath offers users several visualizations designed to evaluate pathway consensus at the database, hierarchy, and gene set level. Users can also opt to directly upload results generated from varying enrichment methods across to visualize variations from these against a set of pathway databases.

### 3.1. Submission form

Once a user has logged into DecoPath, on the Homepage, the input form allows them to upload their files and select parameters to run different analyses or upload results from them **(Figure 2)**. For users opting to run analyses using DecoPath, the workflow depends on the analysis they select. Briefly, GSEA requires the submission of expression datasets, such as from RNA-Seq, microarray, or ChIP-Seq data, accompanied by a design matrix denoting the class labels (e.g., normal and tumor) for samples in the expression dataset. To run ORA, users need only submit a list of genes of interest. For either method, users can select which of the four pathway databases they would like to include in the analysis. By default, genesets from DecoPath which contain merged equivalent pathways are also included in the analysis.

**Figure 2.**
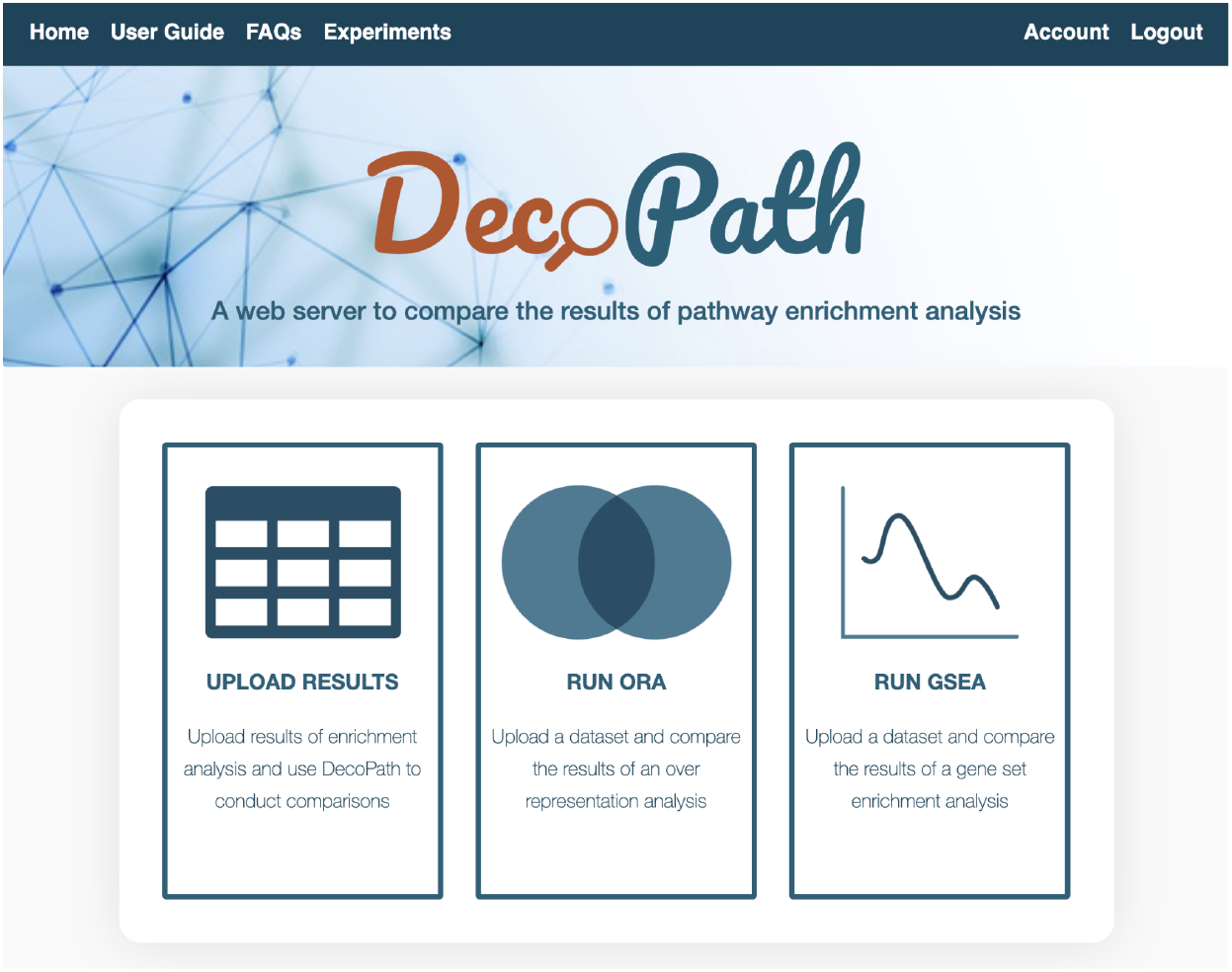
DecoPath homepage. Once a user has logged in, on the homepage, they are provided with the option to either run or submit the results of a pathway analysis. If a user opts to submit the results of an analysis, they can upload their data, select the databases they wish to include, choose the parameter settings for each experiment and optionally perform a concurrent DGE analysis. Once the form has been submitted, users are directed to the Experiments page where they can find visualizations and functionalities to compare and explore the consensus around different pathway databases.

These pathway enrichment methods can also be supplemented by DGE analysis to generate visualizations and identify genes that are differentially expressed according to a fold change cutoff. In order to run DGE analysis, un-normalized read counts in the form of a matrix of integer values is required, as is a design matrix, analogous to the one required for GSEA. For each of these analyses, gene identifiers should be in the form of HUGO Gene Nomenclature Committee (HGNC) symbols. Alternatively, users can opt to download gene set files for pathway databases included in DecoPath, run GSEA, ORA and/or DGE analysis, and upload the results of the analysis to the website. By directly uploading the results, users can also analyze the results of alternative enrichment methods such as Signaling Pathway Impact Analysis (SPIA) (Tarca *et al*., 2008) and EnrichNet (Glaab *et al*., 2012) using DecoPath. Detailed descriptions of the input files can be found in the User Guide and FAQs sections on our website.

### 3.2. Visualizations and analyses

Once users have submitted their query, they are directed to the Experiments page where they can view the status as well as details of their experiments, and explore and visualize their results **(Figure 3)**. To interpret the results of enrichment analysis, we implemented multiple, customized tools intended to provide insights on the consensus across databases, each of which we detail below.

**Figure 3.**
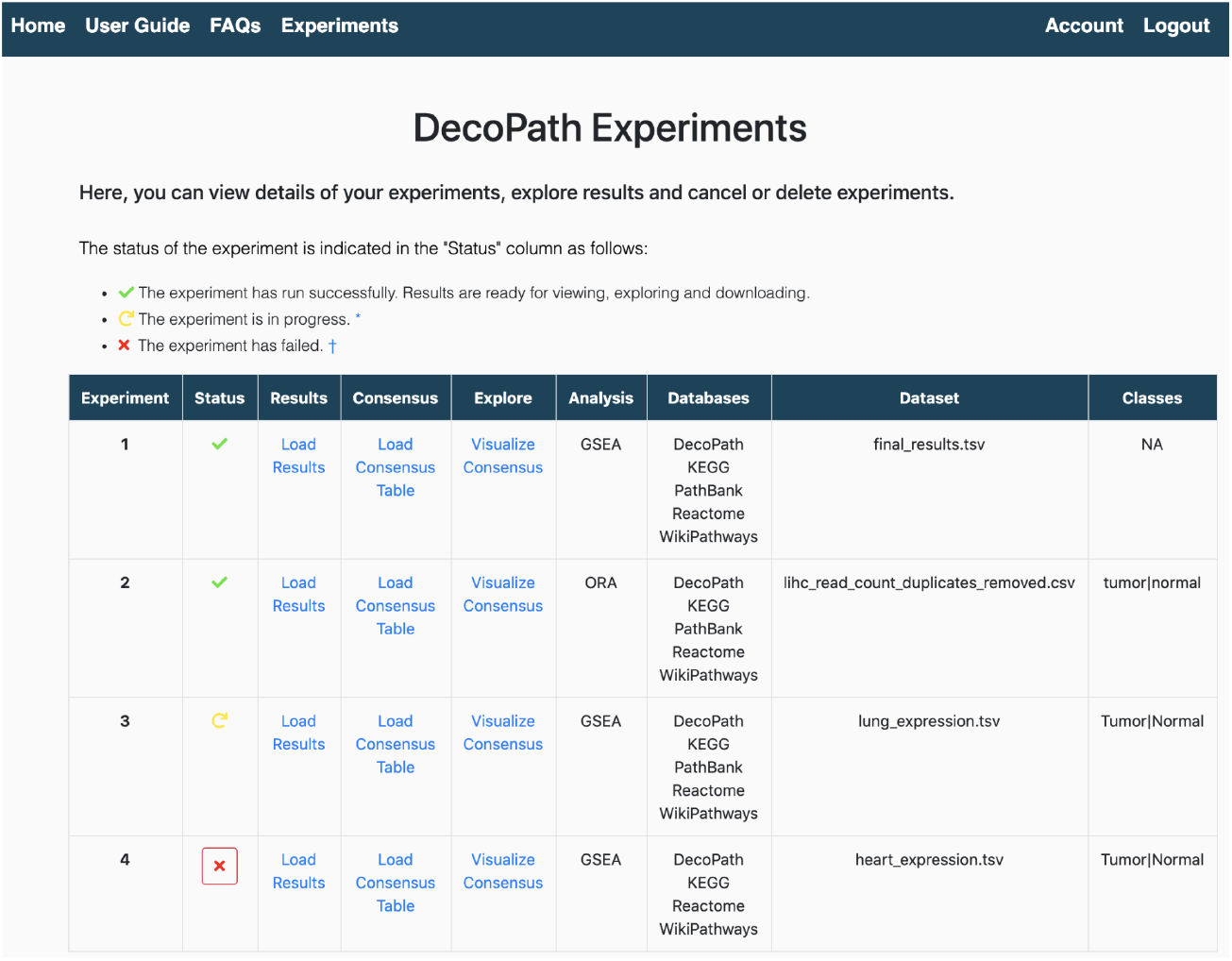
Experiments page. The Experiments page lists details of each of the experiments that were run or uploaded. The status of the experiment is given in the “Status” column, indicating whether the experiment was successfully run, if it is pending or has failed. Through this page, users can then navigate to each of the different visualizations to explore the results of their analysis.

#### 3.2.1. Exploring the consensus across pathway databases

The first visualization summarizes the consensus results of pathway enrichment analysis on multiple databases. For each pathway (row), the table shows the concordance across databases, reflected in terms of the significance value, specifically for ORA, and both the significance value and directionality of the normalized enrichment score (NES) for GSEA **(Figure 4)**. Using this visualization, users can rapidly identify concordant and contradictory pathways and directly compare their results.

**Figure 4.**
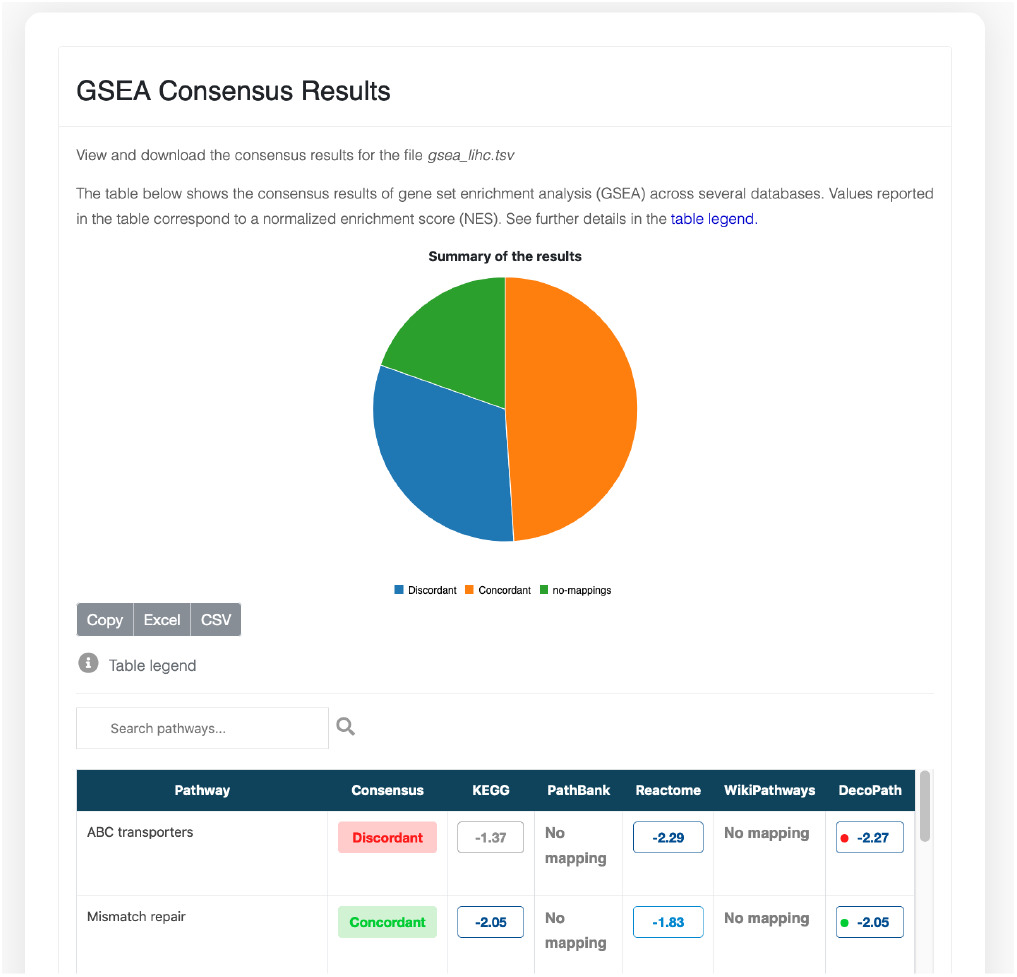
Consensus page. The Consensus page visualization shows the consensus of the results of enrichment analysis across databases at the pathway level. In the case of GSEA, the table displays the NES for a given pathway across each database as well as the NES of the merged gene sets of all equivalent pathways, the latter of which is indicated in the column “DecoPath”. Similarly, ORA results Equivalent pathways mapped enrichment results for equivalent pathways to identify contradictions and discrepancies.

We conducted a case scenario to investigate the results for ORA and GSEA using four pathway databases on the TCGA-LIHC dataset. Among the pathways enriched in ORA which could be found in more than one pathway database, we found 88 concordant pathways (i.e., a given pathway was significantly enriched in the gene list across all databases) and 41 contradictory ones (i.e, a given pathway was significantly enriched in some databases but not in others). Similarly, the results of GSEA revealed 70 concordant and 45 contradictory pathways. Among the contradictory pathways we observed in GSEA, the majority of contradictions pertained to whether or not the pathway was significantly enriched, while 12 pathways also differed in the sign of the NES (i.e., the same pathway was reported as enriched at the top of a ranked gene list for one database and at the bottom for another). Additionally, 53 concordant pathways were common between the results of GSEA and ORA, however, as expected, differences based on the pathway enrichment method were observed. Overall, the results of the LIHC-TCGA dataset for both methods showed that approximately one-third of equivalent pathways were contradictory across the two methods. Thus, the selection of databases, as well as the enrichment method, are important aspects in the experimental design of pathway enrichment analysis. We have observed that the use of one over another can yield discordant results, leading to different interpretations of results depending on the database choice. In the following sections, we illustrate why these results may be discrepant by analyzing the gene sets of a given pathway.

#### 3.2.2. Visualizing consensus through the pathway hierarchy

In the second visualization, users can explore the results of their analysis within the context of a pathway hierarchy **(see Methods)**. This user-friendly and interactive visualization represents the different levels of the pathway hierarchy as circles, each of which represent a child or a parent pathway. In the case of GSEA, pathways that do not show statistically significant (adjusted *p*-value <0.05) differences between groups are coloured gray, while statistically significant ones are coloured red or blue based on the sign of the NES, and shaded by a gradient based on the magnitude of the NES. In the case of ORA, pathways are coloured gray if they are not significant with an adjusted *p*-value < 0.05 and red otherwise. Additionally, the size of the gene sets for each of the pathways is proportional to the size of the circles. Furthermore, interactive visualizations also offer zoom and search functionalities to easily identify pathways of interest. In summary, with this tool, users can not only explore the enrichment results through the entire pathway hierarchy, but also intuitively evaluate equivalent pathways and the size of the pathways, both of which are known to affect results (Karp *et al*, 2021; Mubeen *et al*, 2019).

Continuing the case scenario on the LIHC datasets, this visualization was used to identify major pathways that were enriched in both ORA and GSEA **(Figure 5)**. The organization of pathways into seven major categories allows users to intuitively navigate through the hierarchy and identify pathway groups in which several pathways are enriched. For instance, among all pathways pertaining to metabolism, we observed that lipid and purine metabolism pathways were significantly enriched in both GSEA and ORA, indicating that there was a consensus across both methods and databases. Among other examples of consensus, we found cytokine signaling within the immune system pathways as well as MAP kinase signaling within the signaling pathways significantly enriched in all methods and databases. Finally, contrasting colours of this hierarchical view allow for the rapid identification of contradictory pathways which can then be further analyzed at the gene-level, aided by the following visualization.

**Figure 5.**
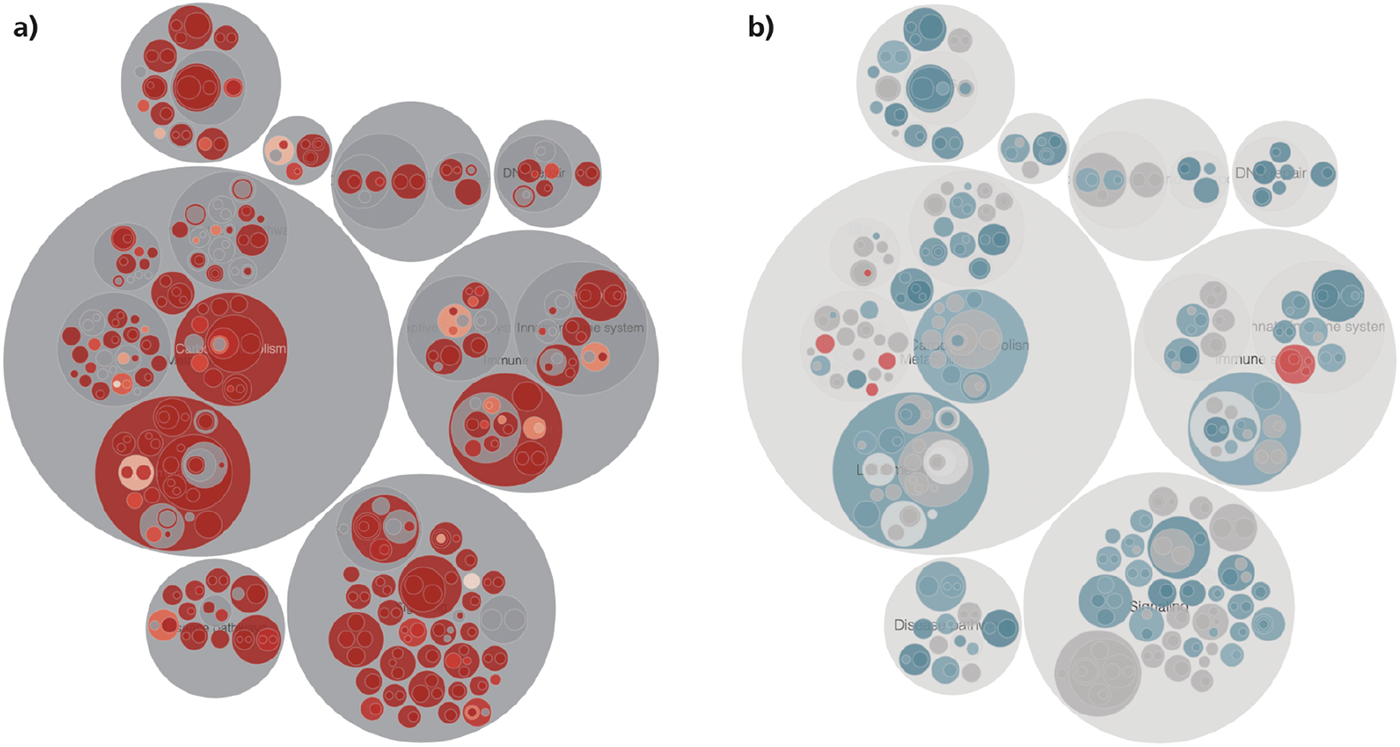
Circle pack visualization of the pathway hierarchy using different pathway enrichment methods. The figure corresponds to the interactive visualizations displaying the results of running ORA **(a)** and GSEA **(b)** on the LIHC dataset. In this visualization, results are customized based on the pathway enrichment method. In the case of Functional Class Scoring (FCS) and Pathway Topology (PT) based methods, the visualization highlights the direction of the dysregulation for each significantly dysregulated pathway as well as for the adjusted *p*-value **(b)**. On the other hand, for ORA, the visualization highlights pathways that are significantly enriched based on an adjusted *p*-value **(a)**.

#### 3.2.3. Analyzing equivalent pathways at the gene level

The third visualization is an interactive Venn diagram that shows the overlap for equivalent pathways at the gene-level. For any given pathway, different databases can disagree in how they define a particular pathway boundary. This can mean that the gene sets of a pathway can differ from database to database. Hence, using different pathway databases can lead to differing results, some of which may be more meaningful than others. In this visualization, we provide a means to analyze exactly which genes may explicate the findings of the pathway analysis. By clicking on the subsets of the Venn diagram, users can display the genes in each of the gene sets. Thus, users can pinpoint the specific genes of the pathway that might contribute to the contradictions observed in the results of the enrichment analysis. If fold changes have additionally been uploaded of differentially expressed genes or DGE analysis has been performed, users can also view the distribution of fold changes of genes in the dataset in an accompanying histogram.

To demonstrate this visualization, we explored both a pathway showing concordant results (i.e., DNA replication pathway) and another showing contradictory results (Pyruvate metabolism) from the results of pathway enrichment on the TCGA-LIHC dataset. In the case of the DNA replication pathway, the results showed that the KEGG, Reactome, and WikiPathways equivalent representations consistently reported NES over 2.0, suggesting that the pathway is regulated in the liver cancer dataset. We then explored the overlap of the gene sets of the DNA replication pathway from the three databases, observing that the log_2_ fold change values for the vast majority of genes in the pathway were positive. As GSEA finds the pathways which are nearest to the top (or bottom) of the ranked list of differentially expressed genes, this can account for the observance of the high NES **(Figure 6, left)**. Similarly, we explored a pathway (i.e., pyruvate metabolism), which had contradictory results in KEGG, Reactome, and PathBank. In this case, these pathway databases disagreed in the direction of regulation of the NES; while the NES of pyruvate metabolism was positive in KEGG and PathBank, the sign of the NES was negative in Reactome. The consensus between KEGG and PathBank is not surprising as the gene sets of the pathway largely overlap **(see Figure 6, right)**, while only 13 of the 31 genes in the Reactome pathway overlap with the other two gene sets. By plotting the distribution of the other 18 genes that are uniquely present in the Reactome pathway, we found that these genes were largely over-expressed, explaining the observed differences in the NES between them. Thus, this example illustrates how this tool can be used to assist in the interpretation of the discrepant results of pathway enrichment analysis.

**Figure 6.**
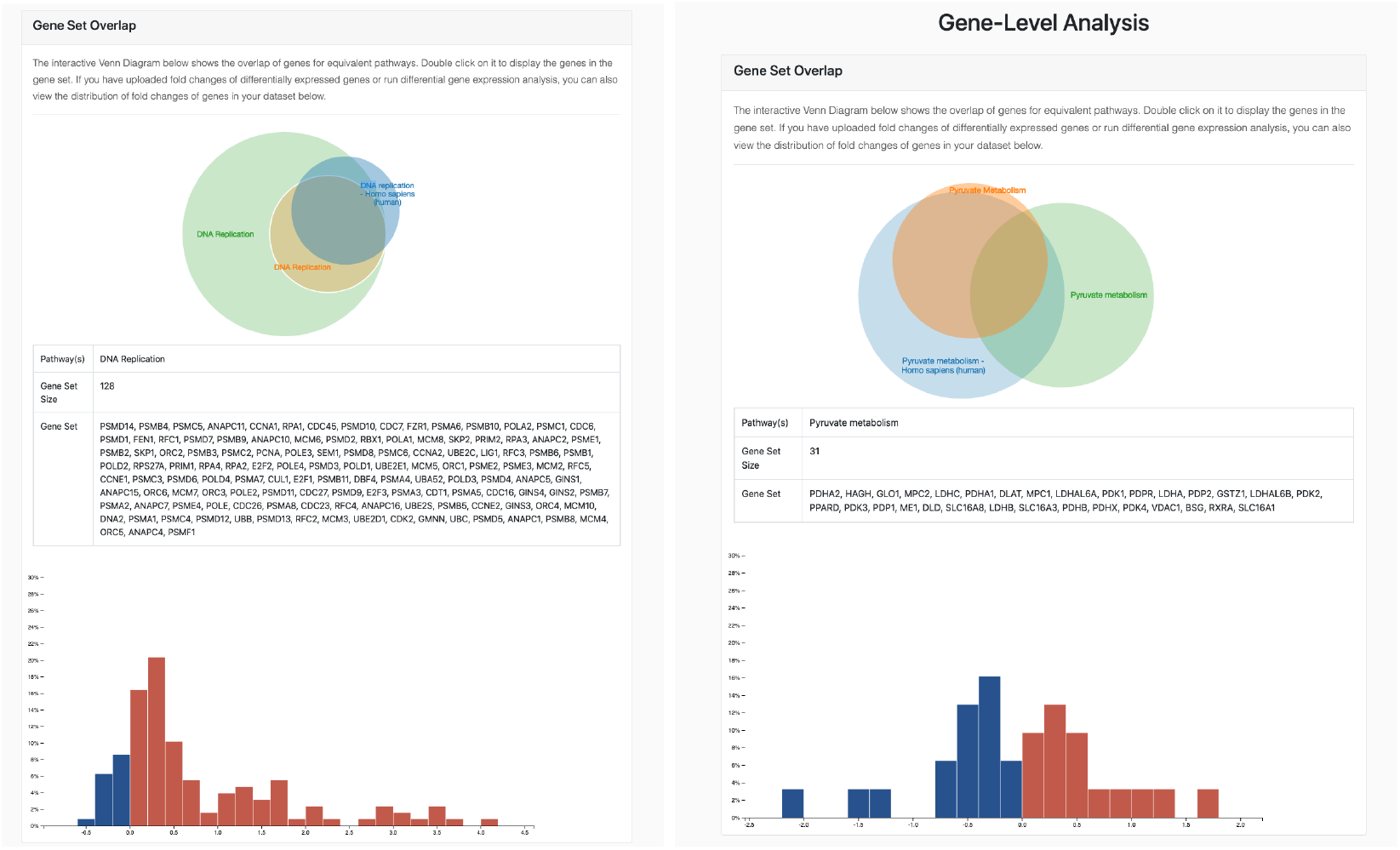
Overlap of gene sets for a given pathway. Venn diagram of the overlap of gene sets for the DNA replication (left) and pyruvate metabolism (right) pathways across multiple databases is shown. By running DGE analysis, users can also view a histogram of the distribution of log_2_ fold changes for differentially expressed genes in their dataset to identify which genes are leading to either consistent or contradictory results of their pathway analysis.

## 4. Discussion

While the popularity of pathway enrichment analysis for the interpretation of -*omics* data has grown over the past two decades and led to the development of over a hundred different methods, recent benchmarks have shown that the selected method can influence results (Geistlinger *et al*., 2020; Nguyen *et al*., 2019; Zyla *et al*., 2019; Mathur *et al*., 2018; Xie *et al*., 2021). Furthermore, the majority of pathway enrichment analyses tend to be conducted on a single pathway database, the choice of which can also impact results of an analysis (Mubeen *et al*., 2019). To date, there have been no tools that allow to directly compare and evaluate the results yielded using different databases or enrichment methods. To address this issue, we have presented DecoPath, the first web application designed to assist in the interpretation of the results of pathway enrichment methods. DecoPath provides users with a broad range of built-in tools and visualization to conduct enrichment analyses and guide them in the interpretation of the results using multiple pathway databases.

Nonetheless, the presented web application is not without its limitations. Firstly, while multiple enrichment methods exist, DecoPath only enables running two of the most popular pathway enrichment analyses. Similarly, DecoPath exclusively contains four pathway databases given the substantial curation effort required to map and harmonize pathway databases. To address these limitations, we enable users to directly upload results from other enrichment methods or pathway mappings from additional databases. Another limitation is the computational power of the server required to run experiments on datasets with a large sample size, or depending on the type of analysis conducted, may not be enough. However, since the source code of the web application is available and DecoPath can be containerized in Docker, users can deploy the web application as per their needs to run more computationally demanding analyses.

In the future, we plan to map and integrate additional databases into DecoPath, as well as more enrichment methods. Furthermore, we would like to develop a more advanced version of the consensus algorithm while taking into account variables such as gene set size and the magnitude of the enrichment score and/or *p*-value. Finally, we hope that our curation effort lays the groundwork for a future overarching pathway ontology with cross-references to databases that could be leveraged and extended by the pathway community.

## Authors’ Contributions

DDF conceived and designed the study. SM implemented the web application and analyzed the data with help from VSB and DDF. YG, SM, and DDF curated the pathway mappings. SM and DDF wrote the paper. ATM, MHA, SM, and DDF acquired the funding.

All authors have read and approved the final manuscript.

## Acknowledgements

We are very grateful to the curators of KEGG, Reactome, WikiPathways, and PathBank for generating the raw content which was used in this work. Furthermore, we would like to thank Vasco Asturiano for developing *circle-packing*, the JavaScript library which is the basis of one of the visualizations of DecoPath.

