## Supplementary File for "DecoPath: A web application for decoding pathway enrichment analysis"

### Supplementary Text 1: Curating a pathway hierarchy

For curating hierarchical pathways, we used two mapping functions: '*equivalentTo*' and '*PartOf*'. *equivalentTo* was used to map two or more analogous pathway representations between the different databases. Furthermore, these equivalent pathways were grouped into 'super pathways' which were identified by a unique identifier. Each equivalent pathway then had a *PartOf* mapping to their corresponding super pathway. Apart from mapping the different pathways, we classified them based on their mechanism of action into 8 major categories with the following prefixes: metabolism pathways (DC1), immune system pathways (DC2), signaling pathways (DC3), communication and transport pathways (DC4), programmed cell death pathways (DC5), disease pathways (DC6), DNA repair and replication pathways (DC7) and others (DC8). Acknowledging the vast classification categories, we sub-classified the major categories DC1 and DC2. Each subcategory was also indexed using hyphenated blocks in the DC prefixed-identifiers to represent their depth in their respective category. Therefore the metabolism pathways (DC1) was further subdivided into amino acid metabolism and derivative pathways (DC1-1), carbon metabolism pathways (DC1-2), lipid metabolism pathways (DC1-3), nucleotide metabolism pathways (DC1-4), vitamin metabolism pathways (DC1-5) and other metabolic pathways (DC1-6). On the other hand, the immune system pathways (DC2) were subdivided into adaptive immune system pathways (DC2-1), innate immune system pathways (DC2-2) and cytokine signalling pathways (DC2-3).

### Supplementary Text 2: Curating pathway mappings between PathBank and each of KEGG, Reactome, and WikiPathways

For mapping of pathways between different databases, we followed the guidelines previously established in Domingo-Fernández *et al.* (2018). First, we identified pathways from PathBank with similar names as any pathway in KEGG, Reactome, or WikiPathways using approximate string matching. When a number of identical strings were captured in this process, a further refinement of the list was done based on manual curation. In this manual curation, two curators compared the description of different pathways in the list to find the potential link between them and the query pathway. Finally, a final scrutiny was conducted on the remaining pathways from PathBank to verify that they did not correspond to any pathway from the other three databases.

### Supplementary Table 1

| GSEA | | ORA | |
|---|---|---|---|
| **Parameter** | **Configuration** | **Parameter** | **Configuration** |
| Maximum gene set size | 1000 | Maximum gene set size | 1000 |
| Minimum gene set size | 15 | Minimum gene set size | 10 |
| Significance threshold | $q$-value < 0.05 | Significance threshold | $q$-value < 0.05 |

| | | | |
|---|---|---|---|
| Method | *signal_to_noise* | - | - |
| Permutation type | phenotype | - | - |
| Number of permutations | 100 | - | - |

**Supplementary Table 1.** Parameter configuration settings for the presented case scenario.
